## Supplemental text for "Evolution of DNA Replication Origin Specification and Gene Silencing Mechanisms"

### Supplementary Information

**Supplementary Method Table 1 | Summary resources and genotype of yeast strains used in this study**

**Supplementary Video 1 | Interaction of the ORC4 α-helix and origin DNA.** The Movie was made with PyMOL software (The PyMOL Molecular Graphics System, Version 2.0 Schrödinger, LLC). F485 and Y486 highlighted in red. R478 and N489 highlighted in orange. V475 and A487 highlighted in brown. The 5' to 3' origin DNA strand is in green color with A/T29, G/T30 logo positions highlighted in Dark purple, while the opposite strand is in grey color with T/A29', C/A30' logo positions highlighted in Light purple.

### Extended Data Figure and Table Legends

**Extended Data Fig. 1 | Orc2 loop exists in species with origin sequence specificity and its interaction with DNA. a,** Multiple sequence alignment of Orc2 among representing eukaryotic species as indicated. Orc2 DNA interacting loop region indicated with species that don't have sequence specific origins shadowed in blue and species that sequence specific origins exist shadowed in pink. **b,** Orc2 AAA+ domain structure superposition among Human Orc2 in teal color (from PDB code 5uj7), *Drosophila* Orc2 in wheat color (from PDB code 4xgc) and *S. cerevisiae* Orc2 in brown color (from PDB code 5udb). Orc2 loop that interact with DNA is colored in red. **c-e,** R390, Y395, W396, and H399 interact with DNA in base-specific (specificity) and base-nonspecific (affinity) manner in ORC-DNA structure at 3Å (PDB code 5zr1). Red asterisks denote the base interaction between amino acid and DNA base. Blue asterisks denote the base-nonspecific interaction between amino acid and DNA phosphate backbone. Prime symbols denote bases on the opposite strand. Bases numbering denote as the positions in logo (see Fig. 2b). **c** shows Orc2 loop origin DNA minor groove insertion with base-specific interaction between W396 and G25, C25' and T26, and base-nonspecific interaction between Y395 and phosphate backbone of T27. **d-e,** same as in **c**, but view in different angles. **d** shows the base-specific interaction between W396 and G25, C25' and T26. **e** shows the base-nonspecific interaction between R390 and phosphate backbone of T23' and base-nonspecific interaction between H399 and phosphate backbone of A24'.

**Extended Data Fig. 2 | Orc4 mutants viability phenotypes in plasmid shuffle assay.** Detailed procedure of plasmid shuffle assay is described in Method, Plasmid shuffle assay. Briefly, strains (*orc4Δ::TRP1* + p*ORC4/URA3* + p*orc4*$^{tested\ allele}$/*LEU2*) were grown overnight in YPD and spotted onto 5-FOA plates with 10-fold serial dilutions starting from $1.5x10^7$ cells and spotted onto YPD plates as control. Mutations were indicated. Strain (*orc4Δ::TRP1* + p*ORC4/URA3* + p*ORC4/LEU2*) and strain (*orc4Δ::TRP1* + p*ORC4/URA3* + p*orc4*$^{null}$/*LEU2*) were spotted as controls. Biological duplicates are denoted as c1 and c2. Plates were cultured under 25˚C, 30˚C, or 37˚C for different days (as indicated) to test their cold or temperature sensitivity. Orc4 mutant phenotypes summarized in Extended Data Table 1.

**Extended Data Fig. 3 | Orc2 mutants viability phenotypes in plasmid shuffle assay.** Detailed procedure of plasmid shuffle assay is described in Method, Plasmid shuffle assay. Briefly, strains (*orc2Δ::TRP1* + p*ORC2/URA3* + p*orc2*$^{tested\ allele}$/*LEU2*) were grown overnight in YPD and spotted onto 5-FOA plates with 10-fold serial dilutions starting from $1.5x10^7$ cells and spotted onto YPD plates as control. Mutations were indicated. Strain (*orc2Δ::TRP1* + p*ORC2/URA3* + p*ORC2/LEU2*) and strain (*orc2Δ::TRP1* + p*ORC2/URA3* + p*orc2*$^{null}$/*LEU2*) were spotted as controls. Biological duplicates are denoted as c1 and c2. Plates were cultured under 25˚C, 30˚C, or 37˚C for different days (as indicated) to test their cold or temperature sensitivity. Orc2 mutant phenotype summarized in Extended Data Table 2.

**Extended Data Fig. 4 | Orc4 protein expression detection and ORC complex formation detection.** Details of method is described in Method, Cell extract preparation, immunoprecipitation, immunoblot analysis and antibodies. NTAP-tagged Orc4 were immunoprecipitated via incubation with IgG beads. Wild type W303 strain, which contains non-tagged Orc4, is used parallelly as a control of pulldown assay. 2% of input and 16.7% of pulled-down lysate were loaded and subsequently immunoblot with anti-Orc4 (SB12) and anti-Orc1 (SB13). Purified ORC complex (including Orc1 and non-tagged Orc4) was also loaded as control for immunoblotting of Orc1 and Orc4. NTAP-tagged Orc4 is around 83kDa (indicated with green arrows) and non-tagged Orc4 is around 56kDa (indicated with yellow arrows), while Orc1 is around 120kDa (indicated with red arrows). Both short and long exposure of blots are indicated.

**Extended Data Fig. 5 | Cell cycle of NTAP-Orc4 integrated strains.** Flow cytometry was done by growing cells into log phase, arresting at G1 phase with α-factor block for 3 hours (around 1~2 cell cycle time length) and then releasing into S phase for different time point (as indicated above on the left). Different time points were harvest and prepared for flow cytometry with method previously described[50]. DNA strained with SYBR green. Orc4 mutants seemed to have hard time going through S phase and progression through mitosis.

**Extended Data Fig. 6 | Viability deficiency phenotype of Orc4 mutants on plasmid can be rescued by integrating Orc4 into genome. a,** Schematic diagram of viability comparison assay between strains surviving dependent on single episomal origin (Orc4 on plasmid, denoted as [P]) or multiple chromosomal origins (Orc4 integrated into genome, denoted as [G]). The [P] strain relies on a CEN-based plasmid with a single replication origin to carry the tested Orc4 mutation and is therefore stringent. **b,** [P] strains (orc4::TRP1 + pORC4/URA3 + porc4/LEU2) and [G] strains (his3::NTAP-orc4$^{mut}$, orc4::TRP1, bar1Δ::TRP1, LEU2::BrdU-Inc + pOrc4/URA3) were grown overnight in YPD and spotted onto 5-FOA plates with 10-fold serial dilutions starting from $1.5x10^7$ cells and spotted onto YPD plates as control. As controls for [P] strains, strain (orc4::TRP1 + pORC4/URA3 + pORC4/LEU2) and strain (orc4::TRP1 + pORC4/URA3 + porc4$^{null}$/LEU2) were spotted. As controls for [G] strains, strain (his3::NTAP-Orc4$^{WT}$, orc4::TRP1, bar1Δ::TRP1, LEU2::BrdU-Inc + pOrc4/URA3) and strain (orc4::TRP1, bar1Δ::TRP1, LEU2::BrdU-Inc + pOrc4/URA3) were spotted. Plates were cultured under 25˚C, 30˚C, or 37˚C for different days (as indicated) to test their temperature sensitivity. The strain lacking a NTAP-tagged Orc4 did not grow on FOA. The viability deficient phenotype of Orc4 mutants on single-origin plasmid seemed to be partially rescued when the mutants are integrated into the genome and survive on multiple origins.

**Extended Data Fig. 7 | ARS motif logos generated from MPOS assay using *ARS317* mutation library. a,** ARS motif logos for Orc4 integrated strains at A and B1 elements generated using mutation library with *ARS317* sequence backbone. Same as Fig 3, top-half of logos representing the origin sequences that were selected-for in MPOS assay and bottom-half of logos representing the origin sequences that were selected-against in MPOS assay. **b,** Magnified view of A element region in a from Orc4$^{WT}$, orc4$^{F485A, Y486A}$, orc4$^{Y486Q}$ strains with bottom-half of logo faded. Dark purple circles indicate the major changes at A/T29, G/T30 logo positions in the Orc4 mutant strains.

**Extended Data Fig. 8 | Principal component analysis and comparison of motif inference methods. a,** PCA analyses of motifs (performed on the *ARS416* library MPOS data, the *ARS317* library MPOS data, or both libraries), and inferred using either information maximization (IM) or enrichment ratios (ER). The variance explained by the first two principal components, corresponding to the x- and y-axes of each plot, is indicated in the upper left corner. The dots within each plot represent biologically independent MPOS experiments, dot color indicates the Orc4 variant assayed, and dot shape indicates the library of mutated ARSs used as input. Note that motifs cluster according to the Orc4 variant assayed, and that this clustering is stronger for information maximization (IM)-inferred motifs compared to enrichment ratio (ER)-inferred motifs. **b,** Total variance across the motifs inferred for each Orc4 variant using either ER or IM inference. IM inference consistently yielded less intra-replicate variance than ER inference (48.3% less on average for *ARS416* motifs and 39.9% less for *ARS317* motifs). This again reflects the robustness of IM inference in the face of experiment-to-experiment variation. **c,** Logos showing the *ARS416* motifs for two Orc4 variants. For clarity, only 20 bp encompassing the essential A element are shown. ER motifs exhibited substantially more variability at key positions than did IM motifs (e.g. rose highlighted positions). Orc4 mutants resulted in consistently and clearly visible differences in the inferred IM motifs (e.g. cyan positions).

**Extended Data Fig. 9 | Replicates of genome-wide replication origin profile. a,** Schematic diagram for genome-wide replication origin profile analysis. Details of method is described in Methods, Genome-wide replication origin profile analysis. Briefly, Yeast cells were $\alpha$-factor blocked in G1 phase for 3 hours and then released into the growth medium (YPD with 200mM HU, 500uM EdU and 0.2mg/ml pronase E) for 90mins before harvest. Flow cytometry was done to check the stage of the cells. DNA is isolated from the harvested cells and sonicated using Bioruptor. EdU labeled newly synthesized DNA is pulldown by Click-iT chemistry with biotinylated azide and Streptavidin T1 magnetic beads. Then Illumina TruSeq Kit is used to establish and amplify the sequencing library. Sequencing data is then analyzed to show peaks on newly synthesized DNA with detailed computational method in Method, Computational analyses of replication origin profile and ChIP-seq data. **b-c,** Replicates of origin firing profiles in Fig. 3. Chromosome IV(ChrIV) is used as representation and replicates are from two independent experiments. **b** shows the direct comparison of two replicates with profiles from $Orc4^{WT}$ and *mrc1*Δ strains shown as examples. **c** shows the genome-wide replication origin firing profiles from the all strains in replicate experiment.

**Extended Data Fig. 10 | Chromatin immuno-precipitation (ChIP) of MCM in Orc4 strains. a-b,** ChIP profile of MCM (anti-Mcm2) in G1 phase ($Orc4^{WT}$, $orc4^{F485I,\ Y486Q}$ and *mrc1*Δ profiles in Fig. 3). Chromosome IV(ChrIV) is used as representation. Genome-wide replication origin firing profiles from Fig. 3 is attached for better reference of origin firing pattern and are shadowed in grey. **a** shows the direct comparison of two independent replicates with ChIP-Mcm2 profiles from $Orc4^{WT}$ and *mrc1*Δ strains shown as examples. **b** shows the ChIP-Mcm2 profiles of the all the strains from one of the two replicates.

**Extended Data Fig. 11 | Genomic origin firing peak heights scatter plot comparisons.** Each dot represent a single replication origin that has its origin firing peak height in $Orc4^{WT}$ (in **b**-**j**) or *mrc1*Δ (in **k**-**t**) as the x-value and its origin firing peak height in $orc4^{mut}$ (in **b**-**j**, **l**-**t**) or $Orc4^{WT}$ (in **k**) as y-value. Two origins exhibited aberrantly large height values, believed to have arisen from read mapping artefacts, and were removed from this analysis. **a**, Illustration diagram for **b**-**j** showing the directions of activation (in green) and repression (in orange) for each replication origin (denote as black dot) in orc4 mutant strains. **b**-**j**, $orc4^{mut}$ strains direct comparison with $Orc4^{WT}$ strain. Height values are in $log_e$ scale. Coefficient of determination values ($R^2$) are shown atop each panel. **k**-**t**, all ten Orc4 strains direct comparison with *mrc1*Δ strain. Origins in **k** with close to zero y-value are early origins, and vice versa. Strains that grow slower generally have small height values indicating that they cannot use the origins efficiently. Therefore, they are forced to use more late origins and have larger origin firing pattern changes as shown in **b**-**j**. Height values are in linear scale.

**Extended Data Fig. 12 | Genomic origin firing peak heights are generally not sequence dependent unless the origin sequence recognition is altered.** DNA sequences under origin firing peaks that were predicted to be ACS were obtained from OriDB[13,37] and used for analysis. MPOS motif scores were assigned by how good the annotated ACSs matched to the MPOS motifs. Correlations between origin firing peak heights (in $\log_{10}$) and MPOS motif scores were evaluated for each annotated ACS. A section indicates the origin peak heights in wild-type strain or wild-type like *orc4* mutant control strain. We took late vs early origin factors into consideration. However, a large value in origin height does not guarantee a high MPOS motif score. B section indicates the orc4 F485 and Y486 mutants that were shown to have their origin sequence recognition altered (Fig. 2c and Fig. 4). C section indicates the orc4 R478 and N489 mutation strains. P-values were also computed to assess the null hypothesis that log EdU heights and motif scores are not correlated; all P-values were Bonferoni corrected (by multiplying by the total number of tests). Significant correlations were indicated: *p<0.05, **p<0.01, ***p<0.001.

**Extended Data Fig. 13 | Non-Y486 *orc4* mutation strains can still efficiently use origins with the "AG" dinucleotide.** Supplemental figure for Fig. 4. Box plots for the 6 orc4 mutant strains that do not have Y486 changed, except for the *orc4*$^{F485Y, Y486F}$ strain that contains a conserved mutation whose strain grows similar to Orc4$^{WT}$ and therefore is an exception of Y486 mutation strain. Y-axis is genomic origin firing peak heights in $\log_{10}$. Each dot denotes an annotated ACS. Box plots elements: the minimum height, first (lower) quartile, median, third (upper) quartile, and maximum height. Diamond denotes outliers that exhibited aberrantly large values.

**Extended Data Table 1 | Summary of plasmid shuffle assay Orc4 mutant phenotypes.**
Orc4 mutant viability phenotypes summarized from plasmid shuffle assay (Extended Data Fig. 1). Indicated symbols denotes different viability phenotypes.

**Extended Data Table 2 | Summary of plasmid shuffle assay Orc2 mutant phenotypes.**
Orc2 mutant viability phenotypes summarized from plasmid shuffle assay (Extended Data Fig. 2). Indicated symbols denotes different viability phenotypes.

**Extended Data Table 3 | Summary of doubling time of NTAP-Orc4 integrated strains.**
Doubling time were calculated based on growth curves (see Fig. 1f) log phase cell concentration.

$$k = \frac{\Delta \log(Cell\ Conc.)}{\Delta Time} = \frac{\log(2)}{Doubling\ Time}$$

$$Doubling\ Time = {\log(2)}/{k}$$

k dictates the slope of linear regression line of growth curves log phase.
