## Supplementary figures and images for "Evolution of DNA Replication Origin Specification and Gene Silencing Mechanisms"

### Supplemental Figure 1

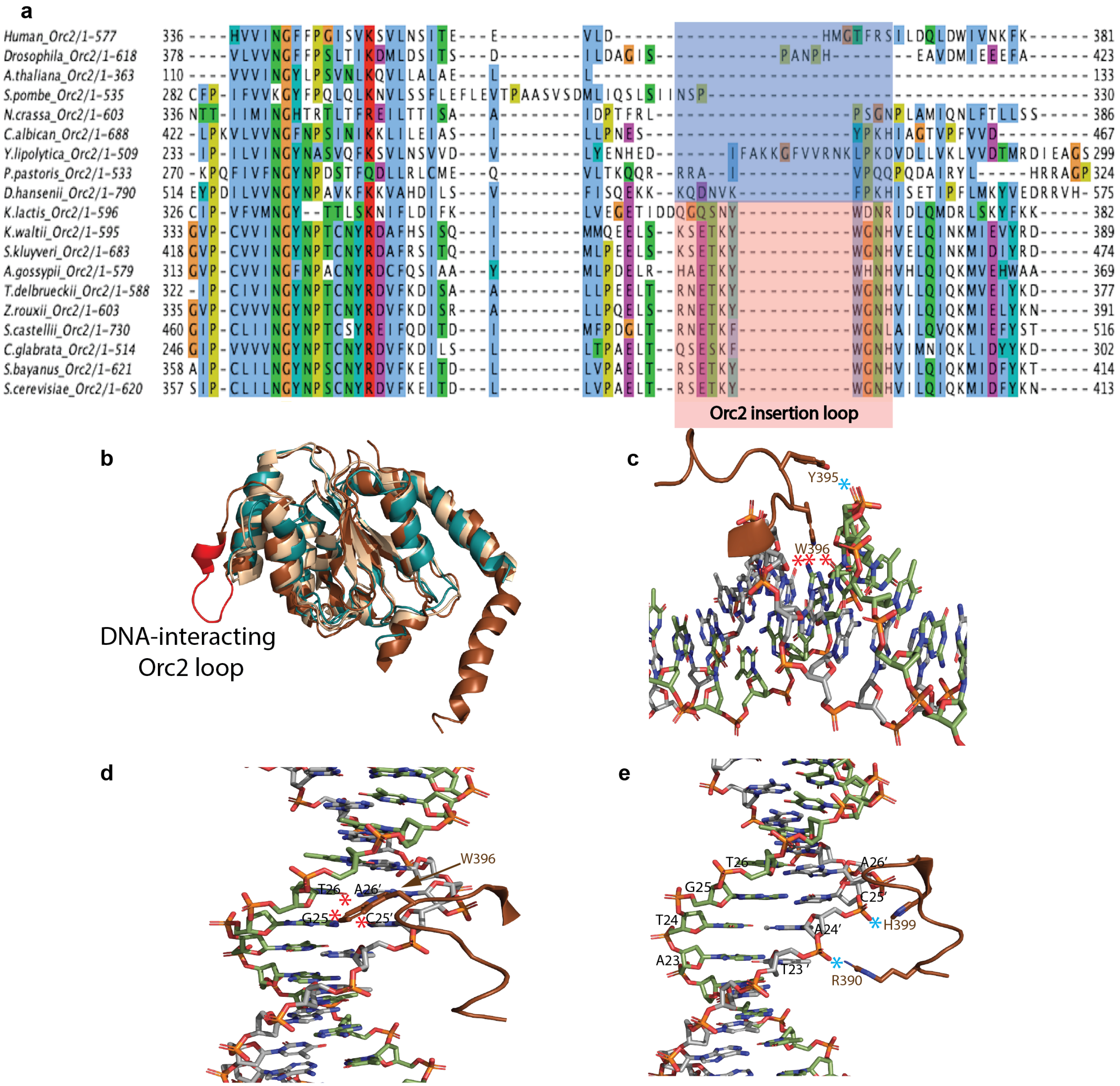


**Extended Data Fig. 1**

### Supplemental Figure 2

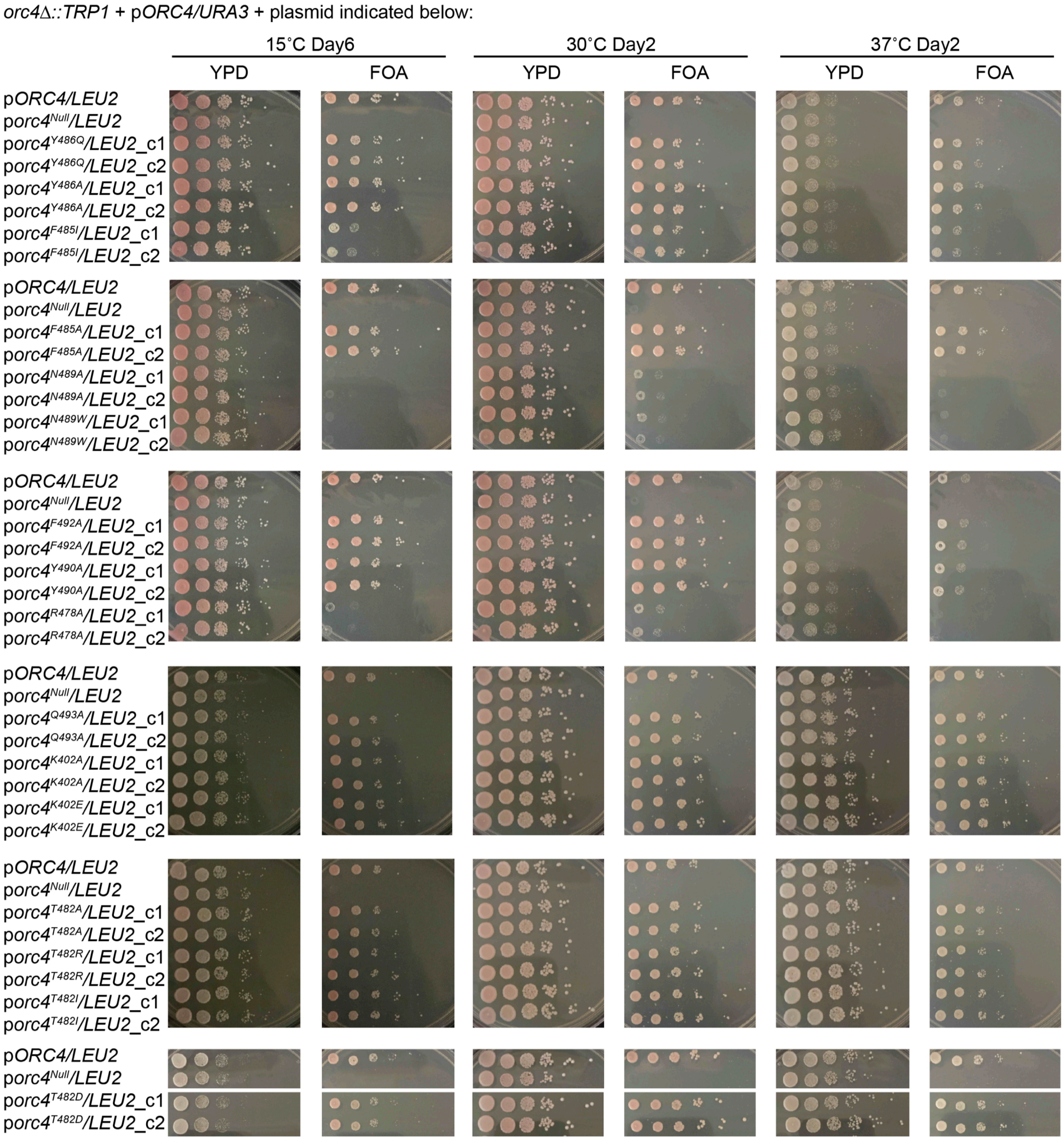


**Extended Data Fig. 2 (1/2)**

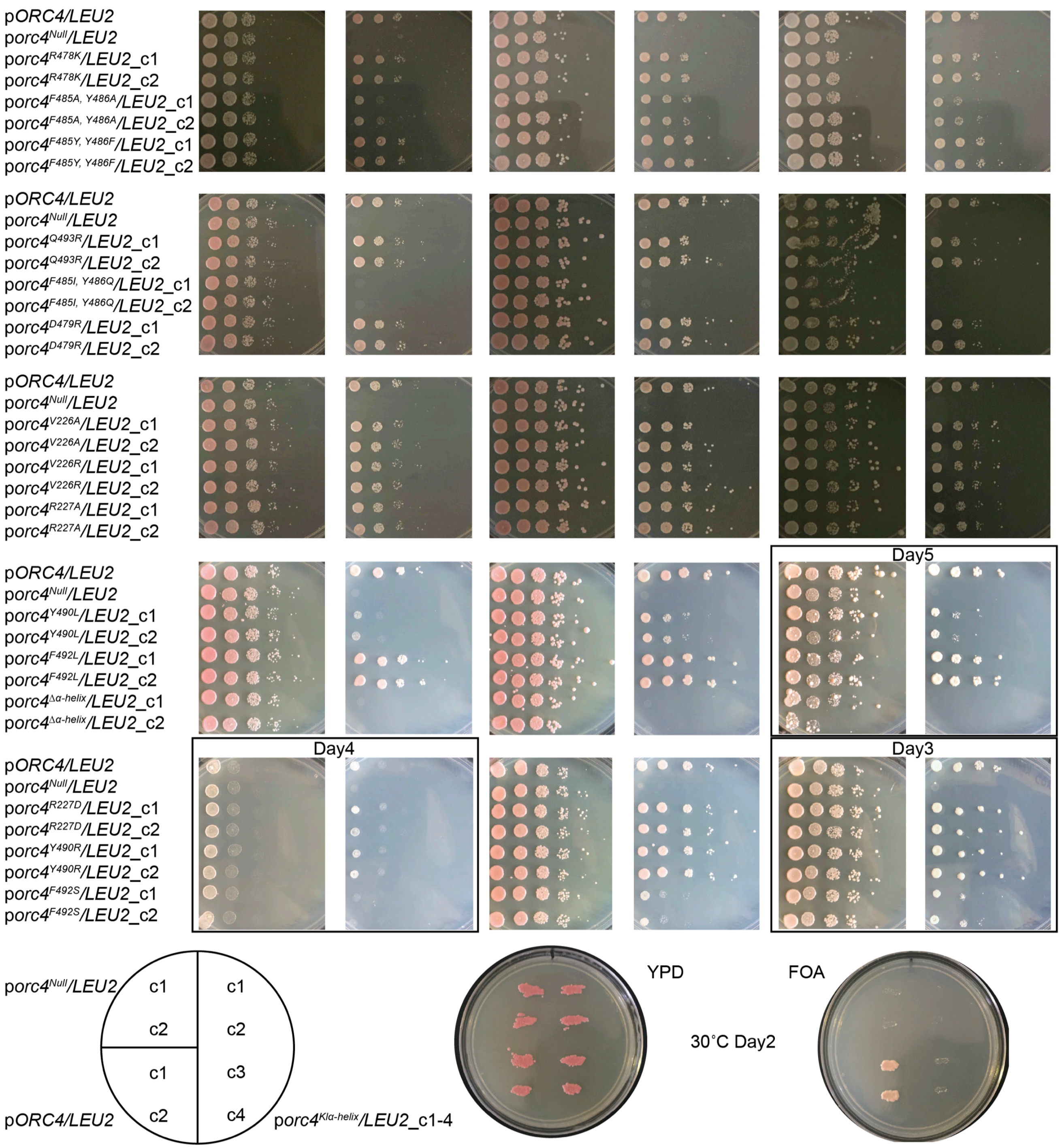


**Extended Data Fig. 2 (2/2)**

### Supplemental Figure 3

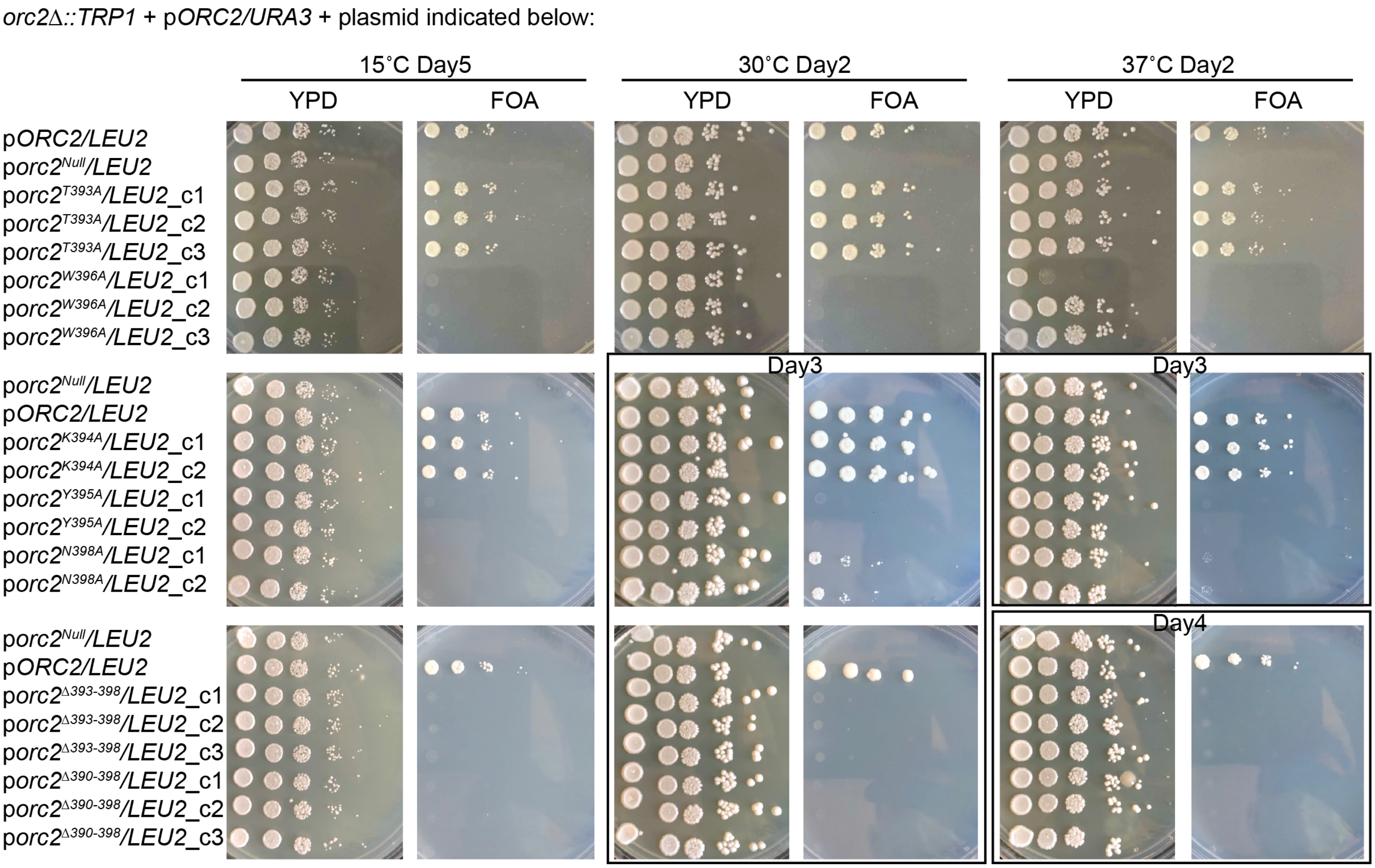


**Extended Data Fig. 3**

### Supplemental Figure 4

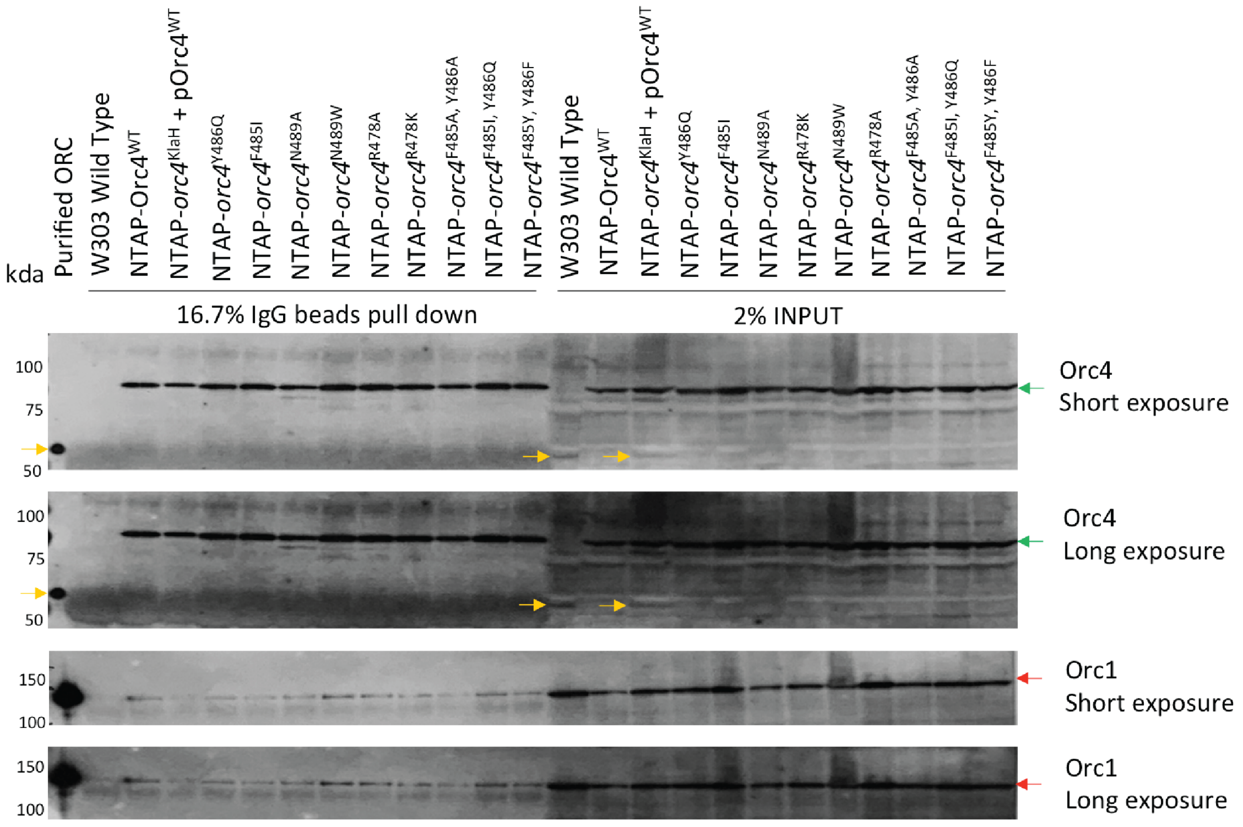


**Extended Data Fig. 4**

### Supplemental Figure 5

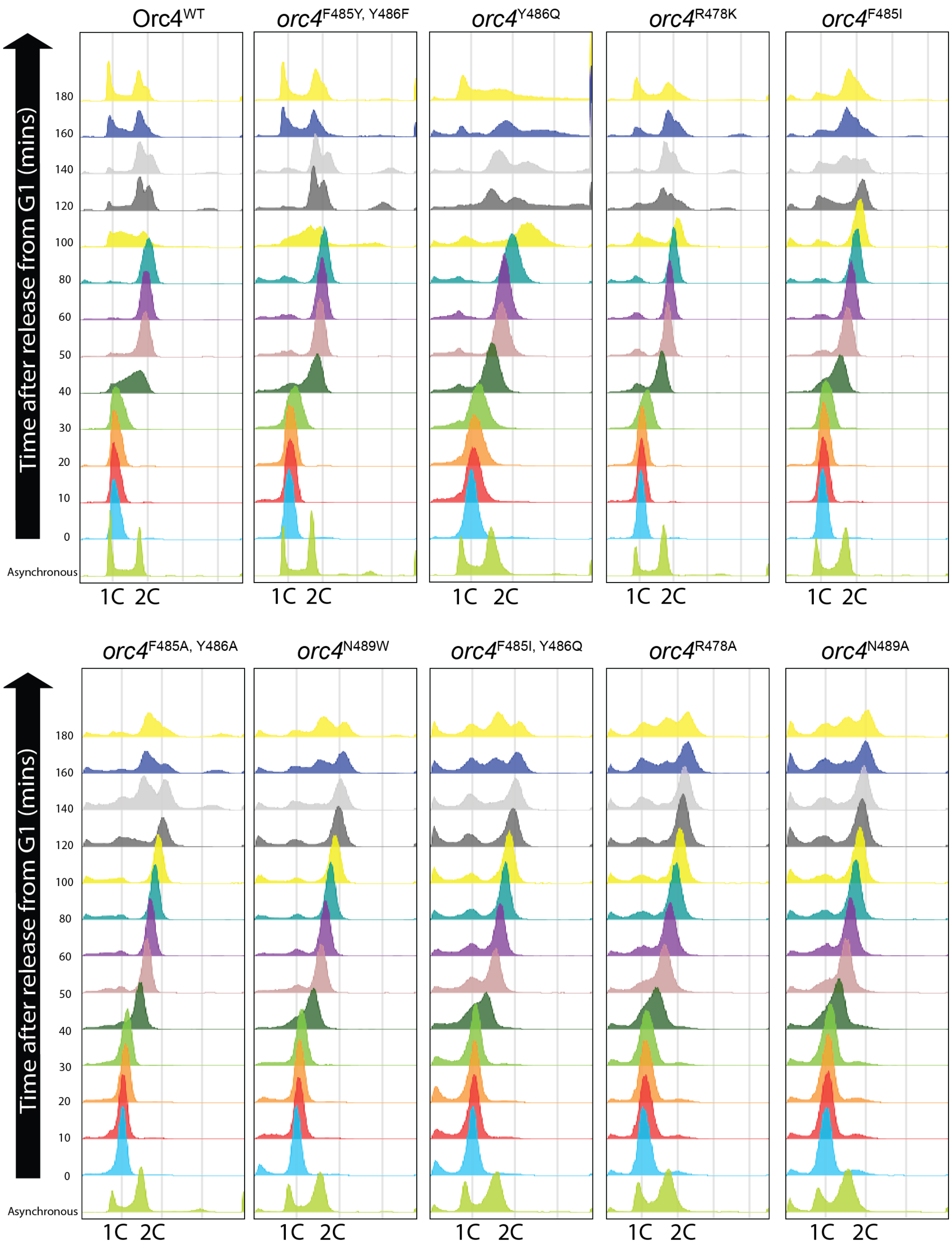


**Extended Data Fig. 5**

### Supplemental Figure 6

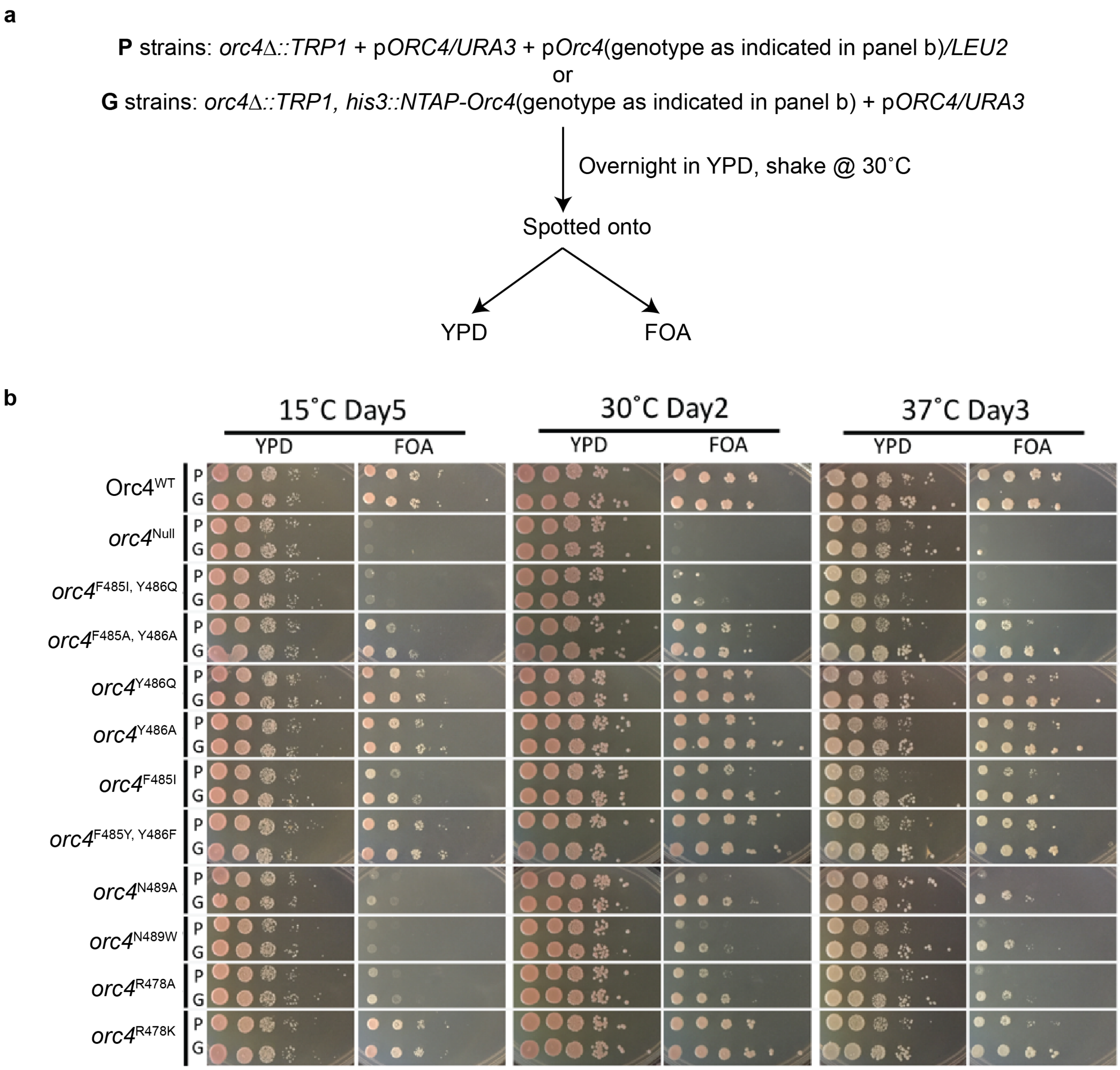


**Extended Data Fig. 6**

### Supplemental Figure 7

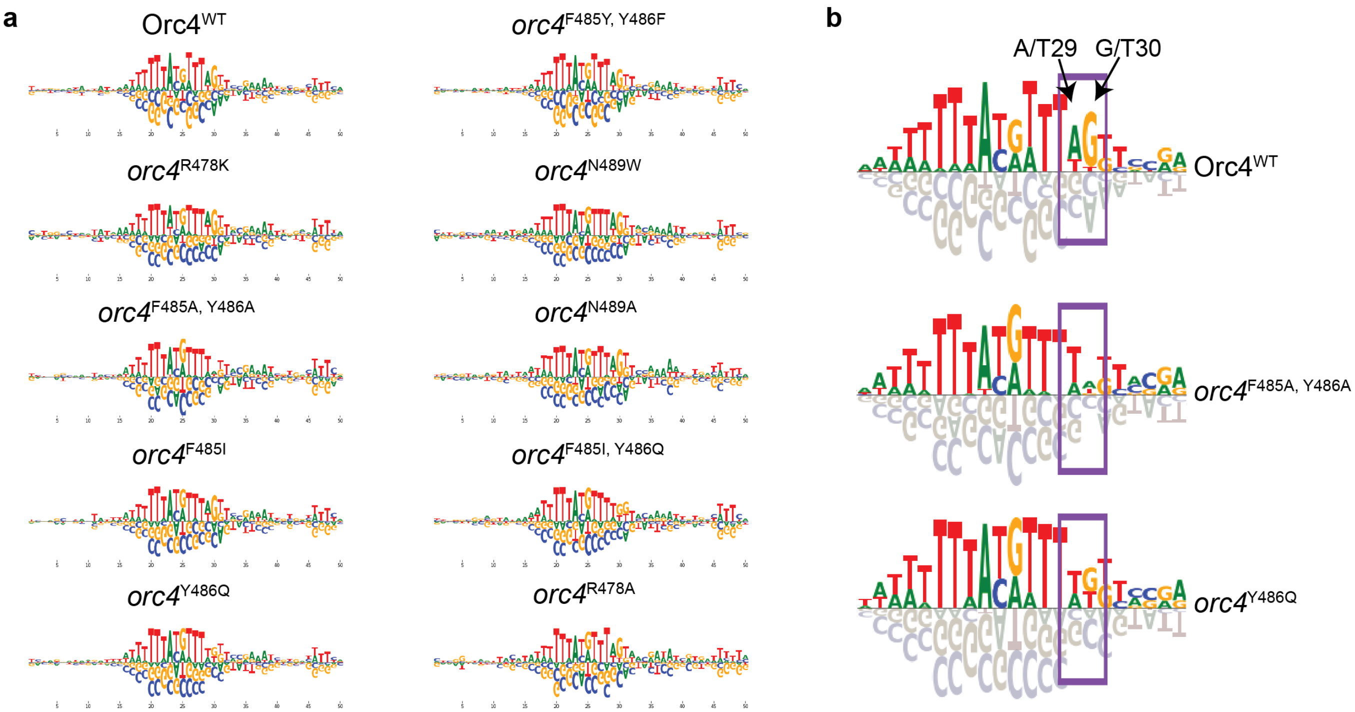


**Extended Data Fig. 7**

### Supplemental Figure 8

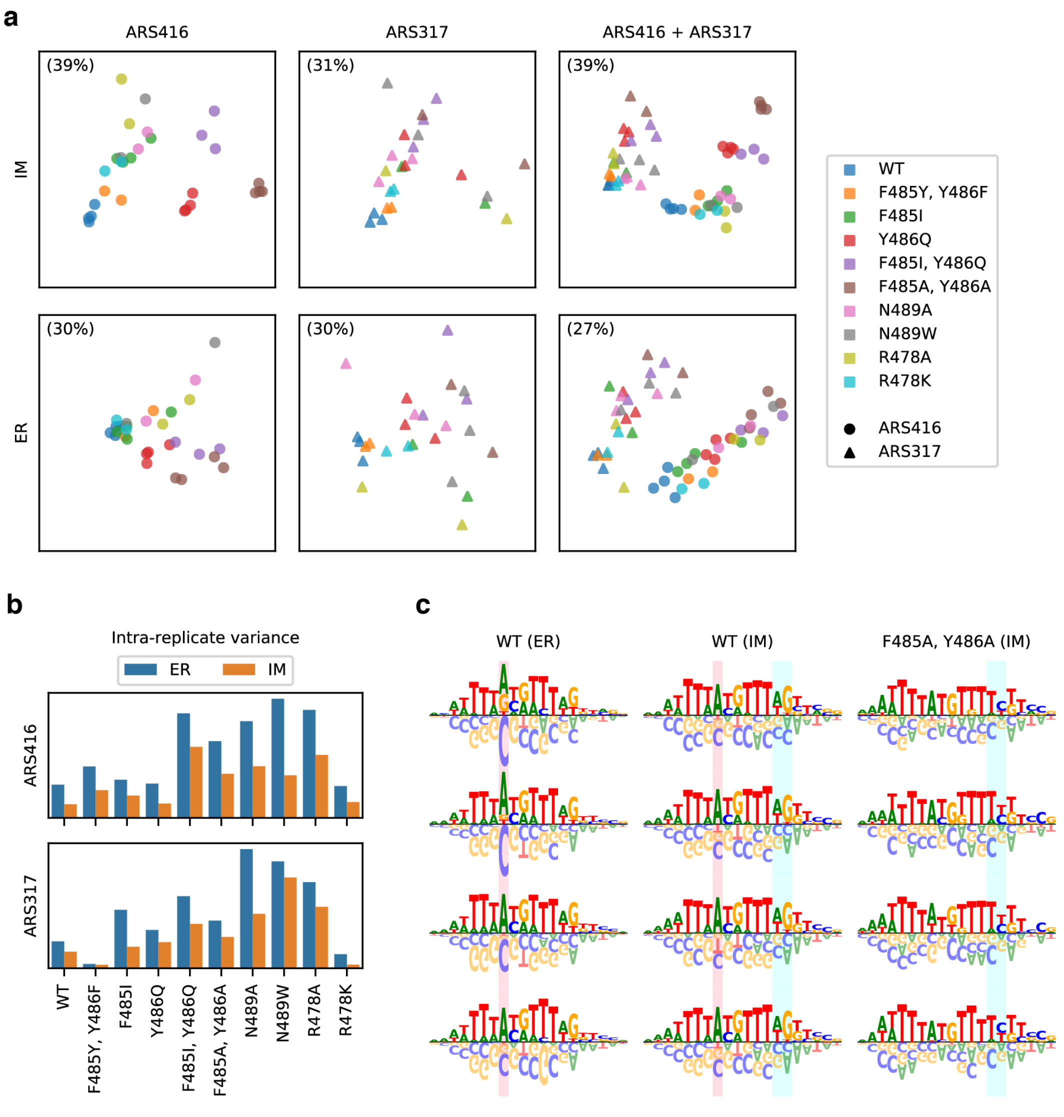


**Extended Data Fig. 8**

### Supplemental Figure 9

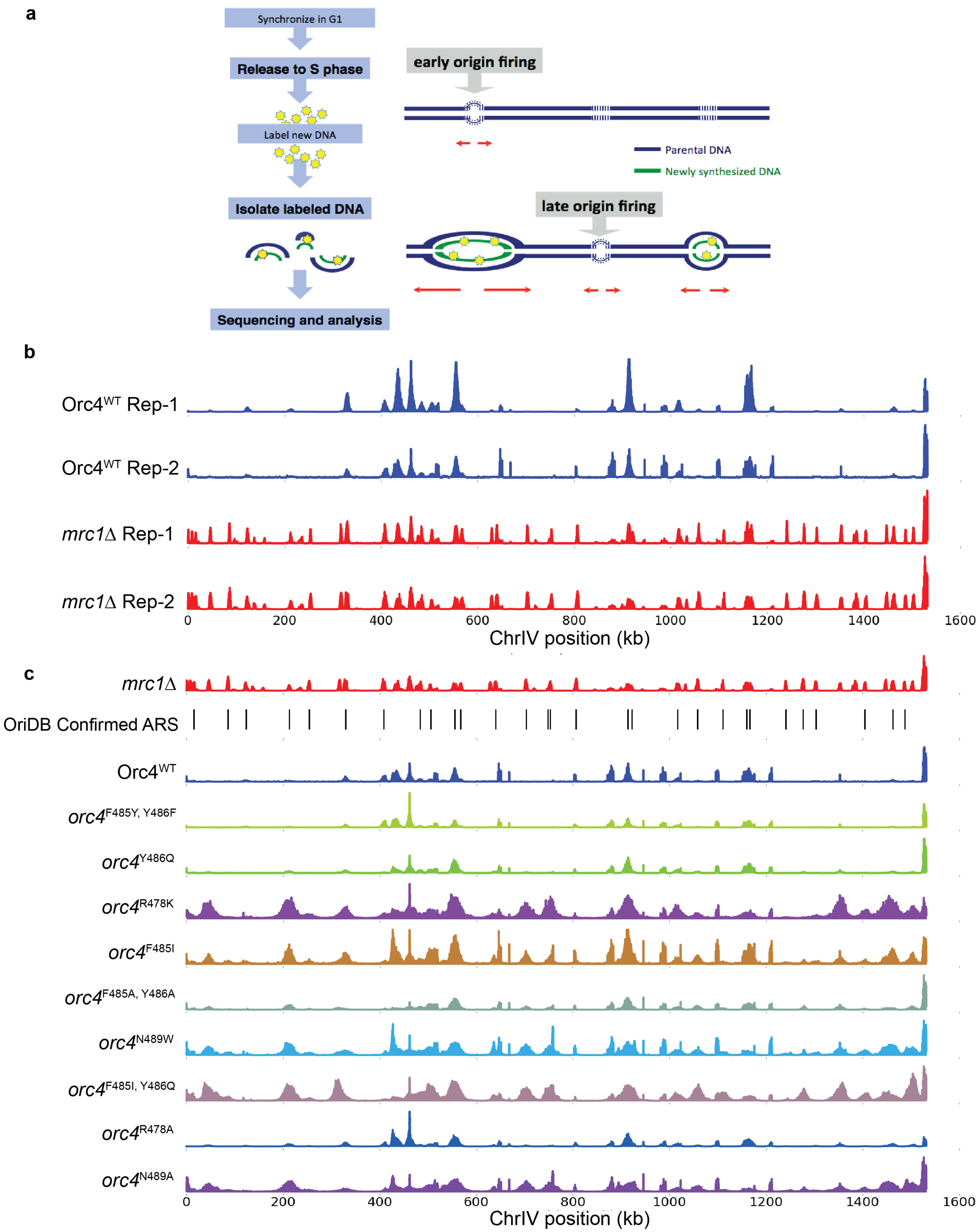


**Extended Data Fig. 9**

### Supplemental Figure 10

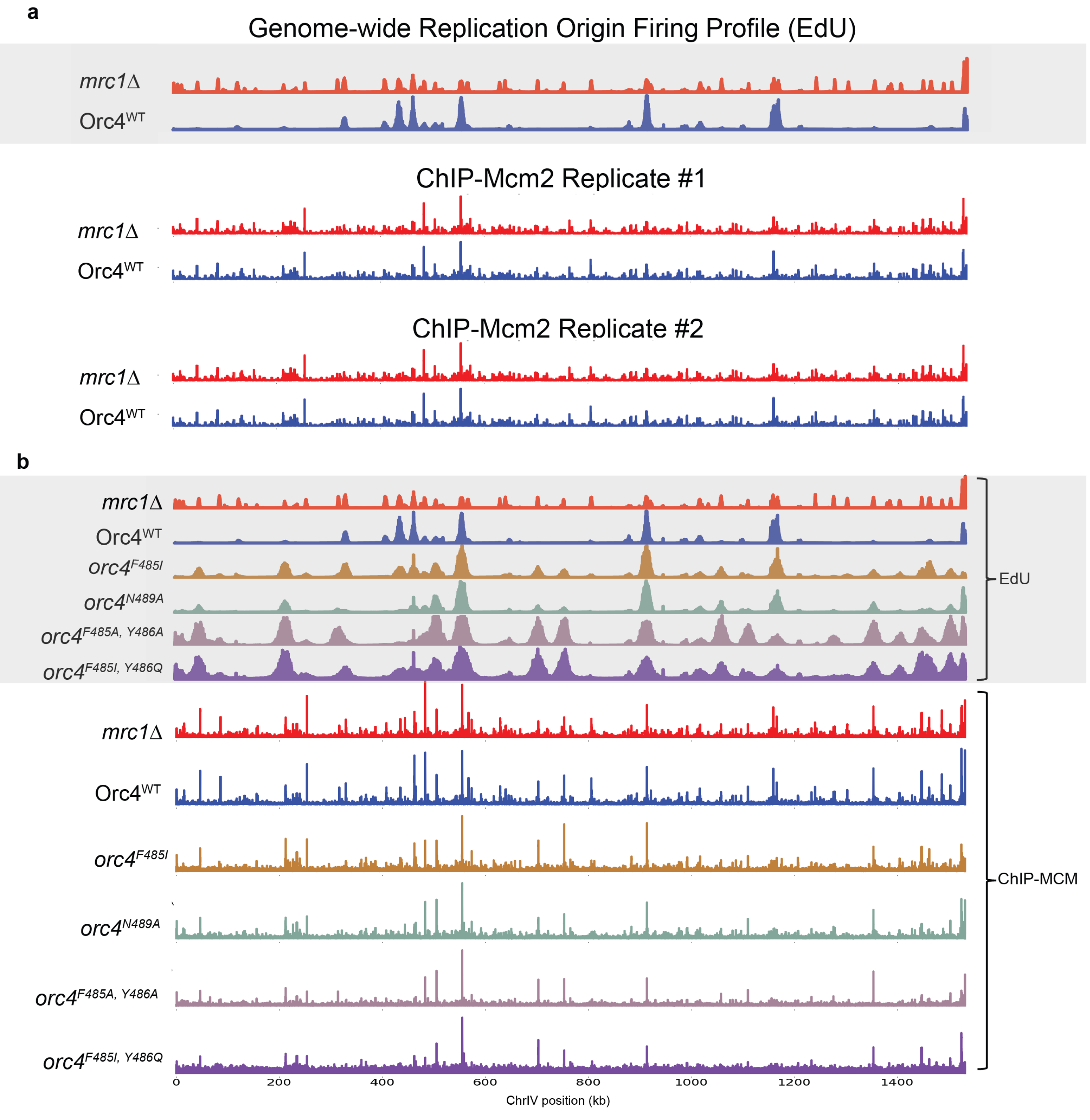


**Extended Data Fig. 10**

### Supplemental Figure 11

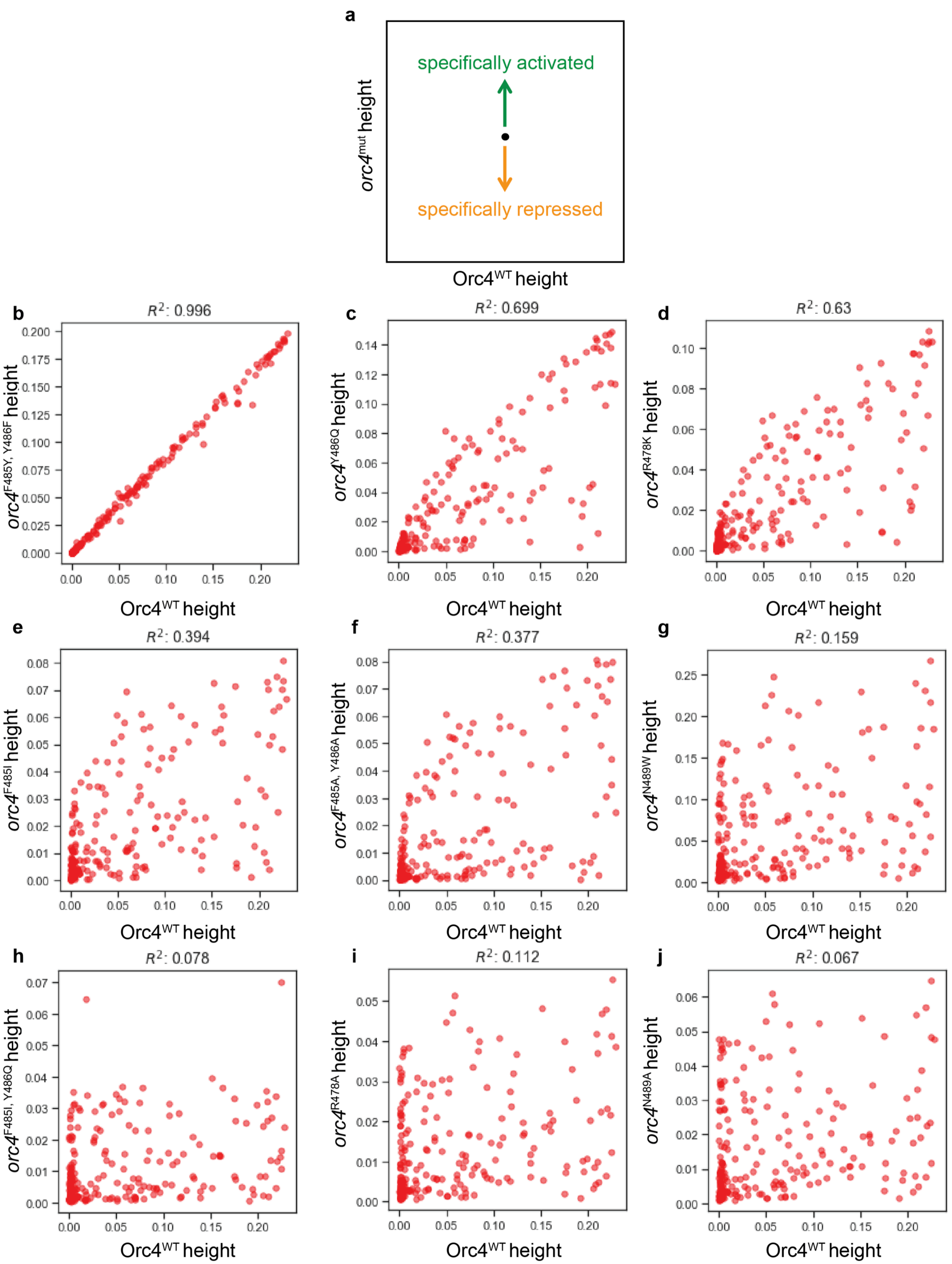


**Extended Data Fig. 11 (1/2)**

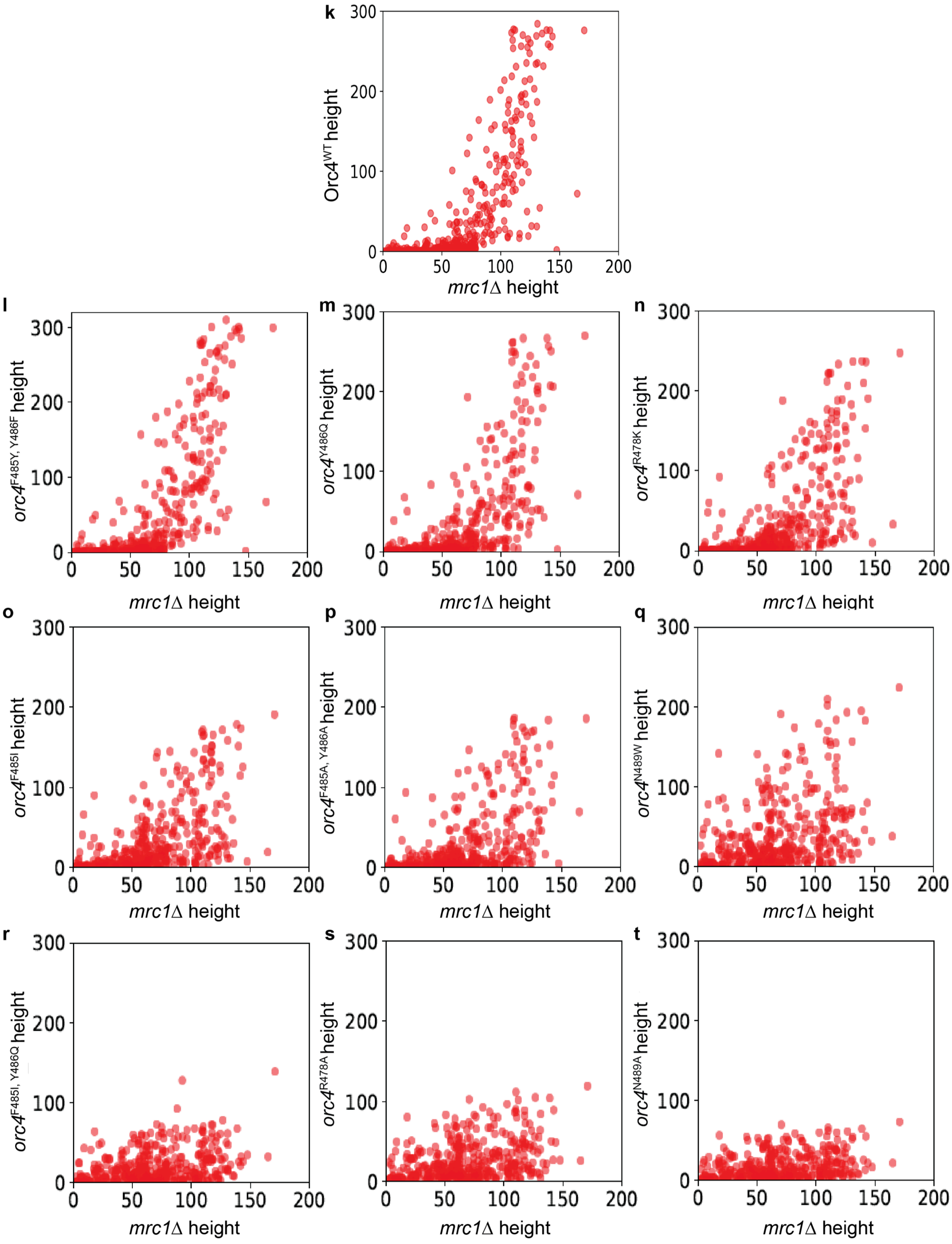


**Extended Data Fig. 11 (2/2)**

### Supplemental Figure 12

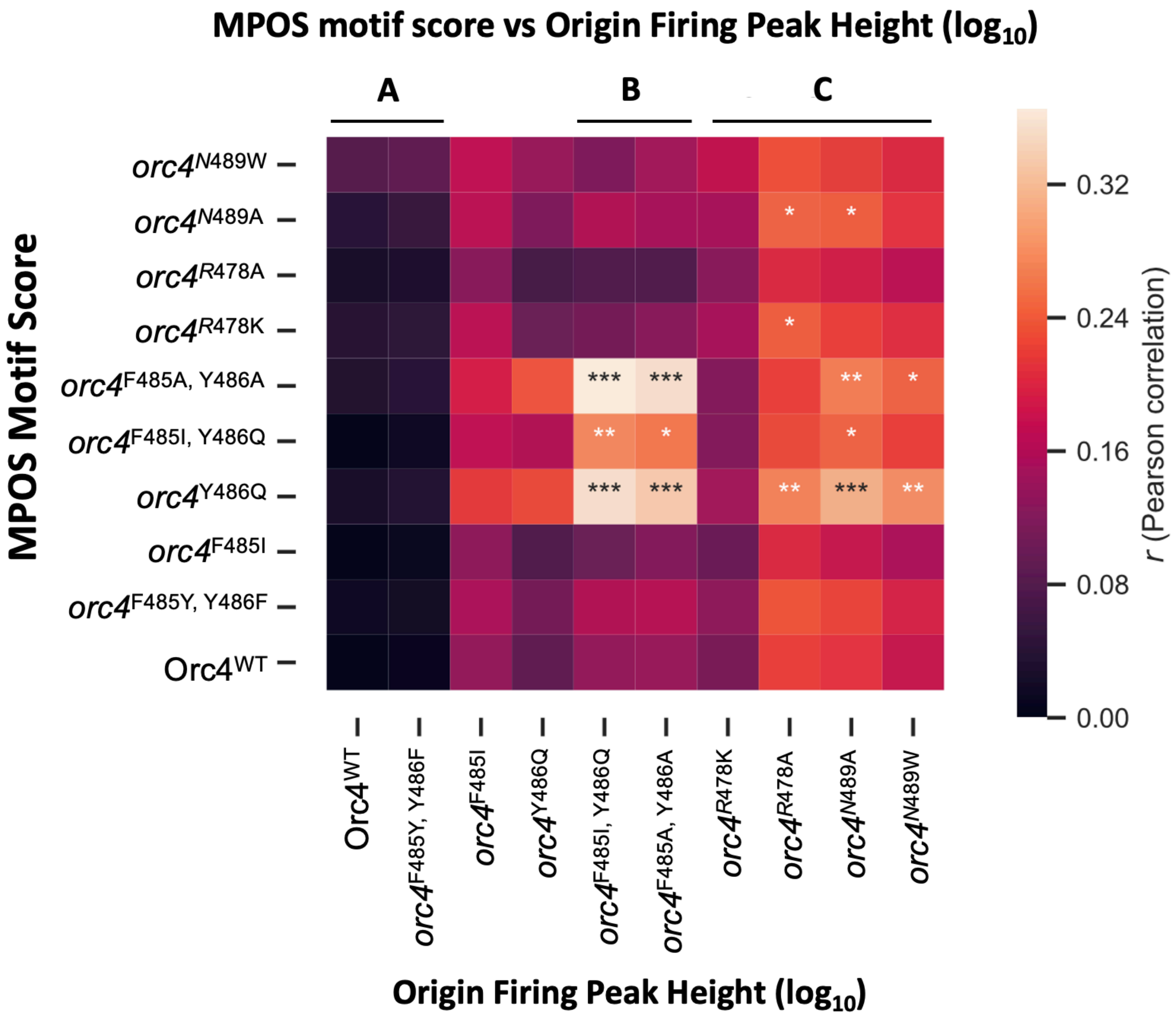


Extended Data Fig. 12

### Supplemental Figure 13

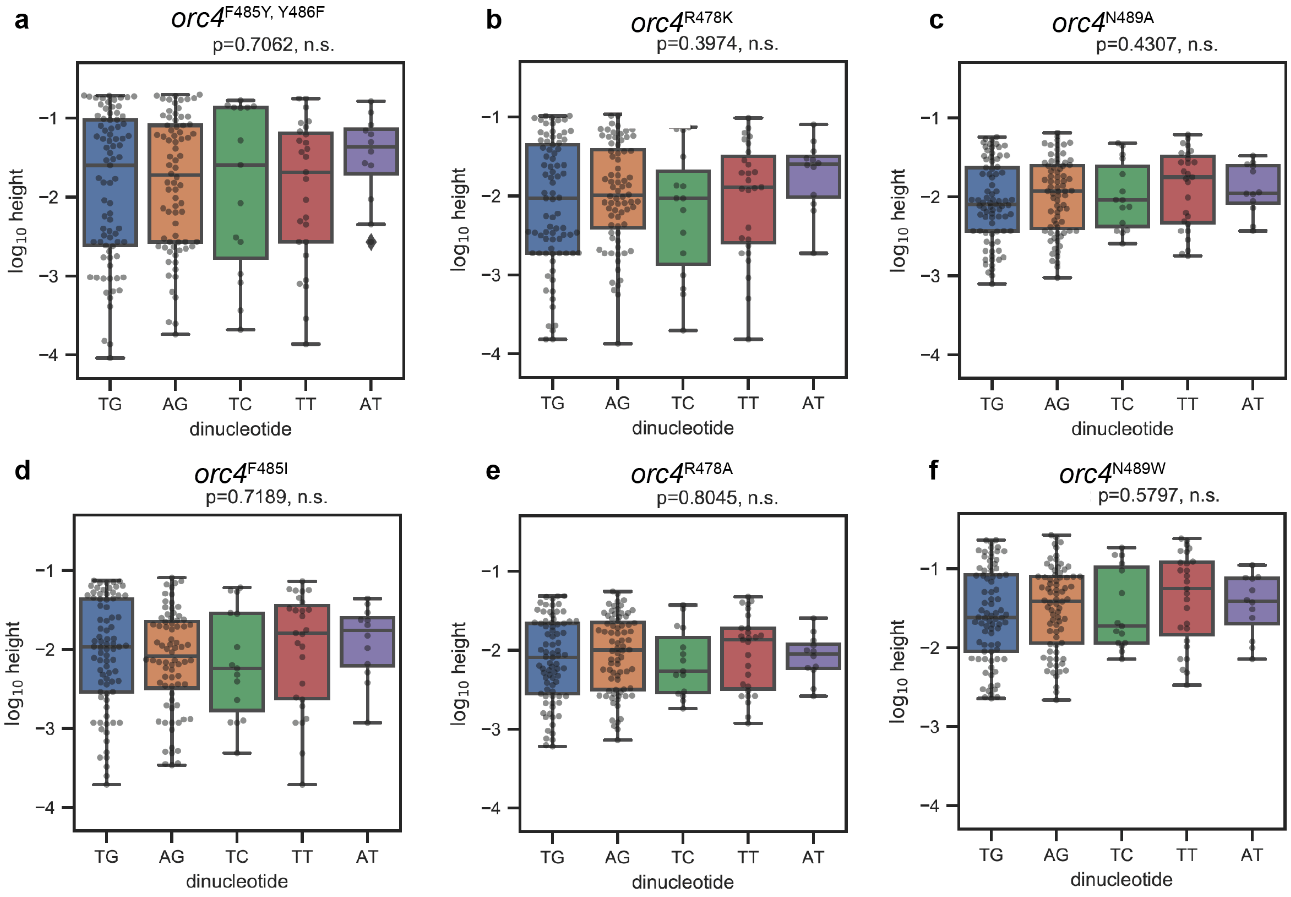


**Extended Data Fig. 13**
