## Supplemental Table 1 for "Evolution of DNA Replication Origin Specification and Gene Silencing Mechanisms"

| Orc4 Mutation | Phenotype |
| --- | --- |
| Klα-helix | # @30˚C (N=4) |
| Δα-helix | # @all three temperature (N=2) |
| F485I, Y486Q | *** @30˚C; # @15˚C (CS); # @37˚C (TS) (N=3) |
| N489W | *** @30˚C; # @15˚C (CS); # @37˚C (TS) (N=3) |
| N489A | *** @30˚C; # @15˚C (CS); # @37˚C (TS) (N=3) |
| R478A | ** @30˚C; *** @15˚C (CS); # @37˚C (TS) (N=3) |
| F485I | * @30˚C; ** @15˚C (CS); * @37˚C (N=3) |
| F485A, Y486A | * @30˚C; ** @15˚C (CS); * @37˚C (N=3) |
| Y490L | * @30˚C; ** @15˚C (CS); * @37˚C (N=2) |
| F492S | * @30˚C; ** @15˚C (CS); * @37˚C (N=2) |
| R478K | - @all three temperature (N=3) |
| Y486Q | - @all three temperature (N=3) |
| F485A | ~ @30˚C; - @15˚C; ~ @37˚C (N=3) |
| Y486A | ~ @30˚C; - @15˚C; ~ @37˚C (N=3) |
| Y490A | ~ @30˚C; - @15˚C; ~ @37˚C (N=3) |
| F492A | ~ @30˚C; - @15˚C; ~ @37˚C (N=3) |
| Q493R | ~ @all three temperature (N=2) |
| D479R | ~ @all three temperature (N=2) |
| V226A | ~ @all three temperature (N=2) |
| V226R | ~ @all three temperature (N=2) |
| R227A | ~ @all three temperature (N=2) |
| Q493A | ~ @all three temperature (N=3) |
| K402A | ~ @all three temperature (N=3) |
| K402E | ~ @all three temperature (N=3) |
| T482D | ~ @all three temperature (N=5) |
| T482R | ~ @all three temperature (N=3) |
| T482A | ~ @all three temperature (N=3) |
| T482I | ~ @all three temperature (N=3) |
| F485Y, Y486F | ~ @all three temperature (N=3) |
| Y490R | ~ @all three temperature (N=3) |
| F492L | ~ @all three temperature (N=3) |
| R227D | ~ @all three temperature (N=3) |

Symbols denotes viability deficient phenotypes. ~ denotes minimal or not deficient; - denotes slightly deficient; * denotes moderate deficient; ** denotes strong deficient; *** denotes severe deficient; # denotes lethal

**Extended Data Table 1**
