## Supplemental Table 2 for "Evolution of DNA Replication Origin Specification and Gene Silencing Mechanisms"

| Orc2 Mutation | Phenotype |
|---|---|
| Δ390-398 | # @all three temperature (N=2) |
| Δ393-398 | # @all three temperature (N=2) |
| W396A | # @all three temperature (N=3) |
| Y395A | # @all three temperature (N=2) |
| N398A | ** @30˚C; # @15˚C (CS); # @37˚C (TS) (N=2) |
| T393A | ~ @all three temperature (N=3) |
| K394A | ~ @all three temperature (N=2) |

Symbols denotes viability deficient phenotypes. ~ denotes minimal or not deficient; - denotes slightly deficient; * denotes moderate deficient; ** denotes strong deficient; *** denotes severe deficient; # denotes lethal

**Extended Data Table 2**
