## Supplemental Table 3 for "Evolution of DNA Replication Origin Specification and Gene Silencing Mechanisms"

| Orc4 Integrated Strains | Doubling Time |
|---|---|
| WT | 87mins |
| F485Y, Y486F | 89mins |
| Y486Q | 91mins |
| R478K | 91.5mins |
| F485I | 93mins |
| F485A, Y486A | 97mins |
| N489W | 112mins |
| F485I, Y486Q | 127mins |
| R478A | 132mins |
| N489A | 134.5mins |

**Extended Data Table 3**
