## Supplemental Methods Table 1 for "Evolution of DNA Replication Origin Specification and Gene Silencing Mechanisms"

| Strain | Genotype | Source |
|---|---|---|
| YB51 | MATα orc4Δ::TRP1 ade2-1 ura3-1 his 3-11, 15 trp1-1 leu2-3,112 can1-100+ pRS415/ORC4 | This lab |
| YS856 | MATa ade2-1 ura3-1 his3-11, 15 trp1-1 leu2-3,112 can1-100 | This study |
| YB410 | MATα orc2Δ::TRP1 ade2-1 ura3-1 his 3-11, 15 trp1-1 leu2-3,112 can1-100+ pRS316/ORC2 | This lab |
| YS3285 | MATa mrc1Δ::KanMX6 bar1Δ::TRP1 URA3::BrdU-Inc ade2-1 ura3-1 his3-11, 15 trp1-1 leu2-3,112 can1-100 | This study |
| YB1519 | MATα orc4Δ::TRP1 ade2-1 ura3-1 his 3-11, 15 trp1-1 leu2-3,112 can1-100+ pRS416/ORC4 + pRS415/ORC4 | This study |
| YB1515 | MATα orc4Δ::TRP1 ade2-1 ura3-1 his 3-11, 15 trp1-1 leu2-3,112 can1-100+ pRS416/ORC4 + pRS415/orc4$^{null}$ | This study |
| YB1639 | MATα orc4Δ::TRP1 ade2-1 ura3-1 his 3-11, 15 trp1-1 leu2-3,112 can1-100+ pRS416/ORC4 + pRS415/orc4$^{\Delta\alpha\text{-}helix}$ | This study |
| YB1514 | MATα orc4Δ::TRP1 ade2-1 ura3-1 his 3-11, 15 trp1-1 leu2-3,112 can1-100+ pRS416/ORC4 + pRS415/orc4$^{Kl\alpha\text{-}helix}$ | This study |
| YB1520 | MATα orc4Δ::TRP1 ade2-1 ura3-1 his 3-11, 15 trp1-1 leu2-3,112 can1-100+ pRS416/ORC4 + pRS415/orc4$^{Y486Q}$ | This study |
| YB1521 | MATα orc4Δ::TRP1 ade2-1 ura3-1 his 3-11, 15 trp1-1 leu2-3,112 can1-100+ pRS416/ORC4 + pRS415/orc4$^{Y486A}$ | This study |
| YB1522 | MATα orc4Δ::TRP1 ade2-1 ura3-1 his 3-11, 15 trp1-1 leu2-3,112 can1-100+ pRS416/ORC4 + pRS415/orc4$^{F485I}$ | This study |
| YB1523 | MATα orc4Δ::TRP1 ade2-1 ura3-1 his 3-11, 15 trp1-1 leu2-3,112 can1-100+ pRS416/ORC4 + pRS415/orc4$^{F485A}$ | This study |
| YB1524 | MATα orc4Δ::TRP1 ade2-1 ura3-1 his 3-11, 15 trp1-1 leu2-3,112 can1-100+ pRS416/ORC4 + pRS415/orc4$^{N489A}$ | This study |
| YB1525 | MATα orc4Δ::TRP1 ade2-1 ura3-1 his 3-11, 15 trp1-1 leu2-3,112 can1-100+ pRS416/ORC4 + pRS415/orc4$^{N489W}$ | This study |
| YB1526 | MATα orc4Δ::TRP1 ade2-1 ura3-1 his 3-11, 15 trp1-1 leu2-3,112 can1-100+ pRS416/ORC4 + pRS415/orc4$^{F492A}$ | This study |
| YB1527 | MATα orc4Δ::TRP1 ade2-1 ura3-1 his 3-11, 15 trp1-1 leu2-3,112 can1-100+ pRS416/ORC4 + pRS415/orc4$^{Y490A}$ | This study |
| YB1528 | MATα orc4Δ::TRP1 ade2-1 ura3-1 his 3-11, 15 trp1-1 leu2-3,112 can1-100+ pRS416/ORC4 + pRS415/orc4$^{R478A}$ | This study |
| YB1555 | MATα orc4Δ::TRP1 ade2-1 ura3-1 his 3-11, 15 trp1-1 leu2-3,112 can1-100+ pRS416/ORC4 + pRS415/orc4$^{Q493A}$ | This study |
| YB1556 | MATα orc4Δ::TRP1 ade2-1 ura3-1 his 3-11, 15 trp1-1 leu2-3,112 can1-100+ pRS416/ORC4 + pRS415/orc4$^{K402A}$ | This study |
| YB1557 | MATα orc4Δ::TRP1 ade2-1 ura3-1 his 3-11, 15 trp1-1 leu2-3,112 can1-100+ pRS416/ORC4 + pRS415/orc4$^{K402E}$ | This study |
| YB1558 | MATα orc4Δ::TRP1 ade2-1 ura3-1 his 3-11, 15 trp1-1 leu2-3,112 can1-100+ pRS416/ORC4 + pRS415/orc4$^{T482A}$ | This study |
| YB1559 | MATα orc4Δ::TRP1 ade2-1 ura3-1 his 3-11, 15 trp1-1 leu2-3,112 can1-100+ pRS416/ORC4 + pRS415/orc4$^{T482R}$ | This study |
| YB1560 | MATα orc4Δ::TRP1 ade2-1 ura3-1 his 3-11, 15 trp1-1 leu2-3,112 can1-100+ pRS416/ORC4 + pRS415/orc4$^{T482I}$ | This study |
| YB1571 | MATα orc4Δ::TRP1 ade2-1 ura3-1 his 3-11, 15 trp1-1 leu2-3,112 can1-100+ pRS416/ORC4 + pRS415/orc4$^{T482D}$ | This study |
| YB1562 | MATα orc4Δ::TRP1 ade2-1 ura3-1 his 3-11, 15 trp1-1 leu2-3,112 can1-100+ pRS416/ORC4 + pRS415/orc4$^{R478K}$ | This study |
| YB1563 | MATα orc4Δ::TRP1 ade2-1 ura3-1 his 3-11, 15 trp1-1 leu2-3,112 can1-100+ pRS416/ORC4 + pRS415/orc4$^{F485A, Y486A}$ | This study |
| YB1564 | MATα orc4Δ::TRP1 ade2-1 ura3-1 his 3-11, 15 trp1-1 leu2-3,112 can1-100+ pRS416/ORC4 + pRS415/orc4$^{F485Y, Y486F}$ | This study |
| YB1635 | MATα orc4Δ::TRP1 ade2-1 ura3-1 his 3-11, 15 trp1-1 leu2-3,112 can1-100+ pRS416/ORC4 + pRS415/orc4$^{F485I, Y486Q}$ | This study |
| YKC01 | MATα orc4Δ::TRP1 ade2-1 ura3-1 his 3-11, 15 trp1-1 leu2-3,112 can1-100+ pRS416/ORC4 + pRS415/orc4$^{F492L}$ | This study |
| YKC02 | MATα orc4Δ::TRP1 ade2-1 ura3-1 his 3-11, 15 trp1-1 leu2-3,112 can1-100+ pRS416/ORC4 + pRS415/orc4$^{Y490L}$ | This study |
| YKC03 | MATα orc4Δ::TRP1 ade2-1 ura3-1 his 3-11, 15 trp1-1 leu2-3,112 can1-100+ pRS416/ORC4 + pRS415/orc4$^{Y490R}$ | This study |
| YKC04 | MATα orc4Δ::TRP1 ade2-1 ura3-1 his 3-11, 15 trp1-1 leu2-3,112 can1-100+ pRS416/ORC4 + pRS415/orc4$^{F492S}$ | This study |
| YB1637 | MATα orc4Δ::TRP1 ade2-1 ura3-1 his 3-11, 15 trp1-1 leu2-3,112 can1-100+ pRS416/ORC4 + pRS415/orc4$^{R227D}$ | This study |
| YEH003 | MATα orc4Δ::TRP1 ade2-1 ura3-1 his 3-11, 15 trp1-1 leu2-3,112 can1-100+ pRS416/ORC4 + pRS415/orc4$^{V226A}$ | This study |
| YEH004 | MATα orc4Δ::TRP1 ade2-1 ura3-1 his 3-11, 15 trp1-1 leu2-3,112 can1-100+ pRS416/ORC4 + pRS415/orc4$^{V226R}$ | This study |
| YEH005 | MATα orc4Δ::TRP1 ade2-1 ura3-1 his 3-11, 15 trp1-1 leu2-3,112 can1-100+ pRS416/ORC4 + pRS415/orc4$^{R227A}$ | This study |
| YEH002 | MATα orc4Δ::TRP1 ade2-1 ura3-1 his 3-11, 15 trp1-1 leu2-3,112 can1-100+ pRS416/ORC4 + pRS415/orc4$^{Q493R}$ | This study |
| YEH001 | MATα orc4Δ::TRP1 ade2-1 ura3-1 his 3-11, 15 trp1-1 leu2-3,112 can1-100+ pRS416/ORC4 + pRS415/orc4$^{D479R}$ | This study |
| YS2251 | MATa bar1Δ::TRP1 ade2-1 ura3-1 his3-11, 15 trp1-1 leu2-3,112 can1-100 | This study |
| YB1549 | MATa bar1Δ::TRP1 LEU2::BrdU-Inc ade2-1 ura3-1 his3-11, 15 trp1-1 leu2-3,112 can1-100 | This study |
| YB1588 | MATa orc4Δ::TRP1 bar1Δ::TRP1 LEU2::BrdU-Inc ade2-1 ura3-1 his3-11, 15 trp1-1 leu2-3,112 can1-100+ pRS416/ORC4 | This study |
| YB1623 | MATa his3::NTAP-ORC4 orc4Δ::TRP1 bar1Δ::TRP1 LEU2::BrdU-Inc ade2-1 ura3-1 his3-11, 15 trp1-1 leu2-3,112 can1-100 + pRS416/ORC4 | This study |
| YB1624 | MATa his3::NTAP-orc4$^{F485I}$ orc4Δ::TRP1 bar1Δ::TRP1 LEU2::BrdU-Inc ade2-1 ura3-1 his3-11, 15 trp1-1 leu2-3,112 can1-100 + pRS416/ORC4 | This study |
| YB1625 | MATa his3::NTAP-orc4$^{N489A}$ orc4Δ::TRP1 bar1Δ::TRP1 LEU2::BrdU-Inc ade2-1 ura3-1 his3-11, 15 trp1-1 leu2-3,112 can1-100+ pRS416/ORC4 | This study |
| YB1630 | MATa his3::NTAP-orc4$^{Kl\alpha\text{-}helix}$ orc4Δ::TRP1 bar1Δ::TRP1 LEU2::BrdU-Inc ade2-1 ura3-1 his3-11, 15 trp1-1 leu2-3,112 can1-100+ pRS416/ORC4 | This study |
| YB1631 | MATa his3::NTAP-orc4$^{F485Y, Y486F}$ orc4Δ::TRP1 bar1Δ::TRP1 LEU2::BrdU-Inc ade2-1 ura3-1 his3-11, 15 trp1-1 leu2-3,112 can1-100 + pRS416/ORC4 | This study |
| YB1632 | MATa his3::NTAP-orc4$^{R478K}$ orc4Δ::TRP1 bar1Δ::TRP1 LEU2::BrdU-Inc ade2-1 ura3-1 his3-11, 15 trp1-1 leu2-3,112 can1-100+ pRS416/ORC4 | This study |
| YB1633 | MATa his3::NTAP-orc4$^{\Delta\alpha\text{-}helix}$ orc4Δ::TRP1 bar1Δ::TRP1 LEU2::BrdU-Inc ade2-1 ura3-1 his3-11, 15 trp1-1 leu2-3,112 can1-100+ pRS416/ORC4 | This study |
| YB1647 | MATa his3::NTAP-orc4$^{N489W}$ orc4Δ::TRP1 bar1Δ::TRP1 LEU2::BrdU-Inc ade2-1 ura3-1 his3-11, 15 trp1-1 leu2-3,112 can1-100+ pRS416/ORC4 | This study |
| YB1648 | MATa his3::NTAP-orc4$^{R478A}$ orc4Δ::TRP1 bar1Δ::TRP1 LEU2::BrdU-Inc ade2-1 ura3-1 his3-11, 15 trp1-1 leu2-3,112 can1-100+ pRS416/ORC4 | This study |
| YB1652 | MATa his3::NTAP-orc4$^{Y486Q}$ orc4Δ::TRP1 bar1Δ::TRP1 LEU2::BrdU-Inc ade2-1 ura3-1 his3-11, 15 trp1-1 leu2-3,112 can1-100+ pRS416/ORC4 | This study |
| YB1653 | MATa his3::NTAP-orc4$^{F485A, Y486A}$ orc4Δ::TRP1 bar1Δ::TRP1 LEU2::BrdU-Inc ade2-1 ura3-1 his3-11, 15 trp1-1 leu2-3,112 can1-100 + pRS416/ORC4 | This study |
| YB1654 | MATa his3::NTAP-orc4$^{F485I, Y486Q}$ orc4Δ::TRP1 bar1Δ::TRP1 LEU2::BrdU-Inc ade2-1 ura3-1 his3-11, 15 trp1-1 leu2-3,112 can1-100+ pRS416/ORC4 | This study |
| YB1649 | MATa his3::NTAP-ORC4 orc4Δ::TRP1 bar1Δ::TRP1 LEU2::BrdU-Inc ade2-1 ura3-1 his3-11, 15 trp1-1 leu2-3,112 can1-100 | This study |
| YB1650 | MATa his3::NTAP-orc4$^{F485I}$ orc4Δ::TRP1 bar1Δ::TRP1 LEU2::BrdU-Inc ade2-1 ura3-1 his3-11, 15 trp1-1 leu2-3,112 can1-100 | This study |
| YB1655 | MATa his3::NTAP-orc4$^{N489A}$ orc4Δ::TRP1 bar1Δ::TRP1 LEU2::BrdU-Inc ade2-1 ura3-1 his3-11, 15 trp1-1 leu2-3,112 can1-100 | This study |
| YB1656 | MATa his3::NTAP-orc4$^{F485Y, Y486F}$ orc4Δ::TRP1 bar1Δ::TRP1 LEU2::BrdU-Inc ade2-1 ura3-1 his3-11, 15 trp1-1 leu2-3,112 can1-100 | This study |
| YB1657 | MATa his3::NTAP-orc4$^{R478K}$ orc4Δ::TRP1 bar1Δ::TRP1 LEU2::BrdU-Inc ade2-1 ura3-1 his3-11, 15 trp1-1 leu2-3,112 can1-100 | This study |
| YB1658 | MATa his3::NTAP-orc4$^{N489W}$ orc4Δ::TRP1 bar1Δ::TRP1 LEU2::BrdU-Inc ade2-1 ura3-1 his3-11, 15 trp1-1 leu2-3,112 can1-100 | This study |
| YB1659 | MATa his3::NTAP-orc4$^{R478A}$ orc4Δ::TRP1 bar1Δ::TRP1 LEU2::BrdU-Inc ade2-1 ura3-1 his3-11, 15 trp1-1 leu2-3,112 can1-100 | This study |
| YB1660 | MATa his3::NTAP-orc4$^{Y486Q}$ orc4Δ::TRP1 bar1Δ::TRP1 LEU2::BrdU-Inc ade2-1 ura3-1 his3-11, 15 trp1-1 leu2-3,112 can1-100 | This study |
| YB1661 | MATa his3::NTAP-orc4$^{F485A, Y486A}$ orc4Δ::TRP1 bar1Δ::TRP1 LEU2::BrdU-Inc ade2-1 ura3-1 his3-11, 15 trp1-1 leu2-3,112 can1-100 | This study |
| YB1662 | MATa his3::NTAP-orc4$^{F485I, Y486Q}$ orc4Δ::TRP1 bar1Δ::TRP1 LEU2::BrdU-Inc ade2-1 ura3-1 his3-11, 15 trp1-1 leu2-3,112 can1-100 | This study |
| YB1620 | MATα orc2Δ::TRP1 ade2-1 ura3-1 his 3-11, 15 trp1-1 leu2-3,112 can1-100+ pRS316/ORC2 + pRS415/orc2$^{null}$ | This study |
| YB1619 | MATα orc2Δ::TRP1 ade2-1 ura3-1 his 3-11, 15 trp1-1 leu2-3,112 can1-100+ pRS316/ORC2 + pRS415/ORC2 | This study |
| YB1621 | MATα orc2Δ::TRP1 ade2-1 ura3-1 his 3-11, 15 trp1-1 leu2-3,112 can1-100+ pRS316/ORC2 + pRS415/orc2$^{T393A}$ | This study |
| YB1611 | MATα orc2Δ::TRP1 ade2-1 ura3-1 his 3-11, 15 trp1-1 leu2-3,112 can1-100+ pRS316/ORC2 + pRS415/orc2$^{W396A}$ | This study |
| YKC05 | MATα orc2Δ::TRP1 ade2-1 ura3-1 his 3-11, 15 trp1-1 leu2-3,112 can1-100+ pRS316/ORC2 + pRS415/orc2$^{K394A}$ | This study |
| YKC06 | MATα orc2Δ::TRP1 ade2-1 ura3-1 his 3-11, 15 trp1-1 leu2-3,112 can1-100+ pRS316/ORC2 + pRS415/orc2$^{Y395A}$ | This study |
| YKC07 | MATα orc2Δ::TRP1 ade2-1 ura3-1 his 3-11, 15 trp1-1 leu2-3,112 can1-100+ pRS316/ORC2 + pRS415/orc2$^{N398A}$ | This study |
| YKC08 | MATα orc2Δ::TRP1 ade2-1 ura3-1 his 3-11, 15 trp1-1 leu2-3,112 can1-100+ pRS316/ORC2 + pRS415/orc2$^{\Delta 393\text{-}398}$ | This study |
| YKC09 | MATα orc2Δ::TRP1 ade2-1 ura3-1 his 3-11, 15 trp1-1 leu2-3,112 can1-100+ pRS316/ORC2 + pRS415/orc2$^{\Delta 390\text{-}398}$ | This study |

**Supplementary Method Table 1**
